# Activin B Promotes the Initiation and Progression of Liver Fibrosis

**DOI:** 10.1101/2021.04.27.441623

**Authors:** Yan Wang, Matthew Hamang, Alexander Culver, Huaizhou Jiang, Praveen Kusumanchi, Naga Chalasani, Suthat Liangpunsakul, Benjamin C. Yaden, Guoli Dai

**Author notes:** **Corresponding Authors:** Benjamin Yaden, PhD, Adjunct Professor, Department of Biology, Indiana University-Purdue University Indianapolis, Indianapolis, IN, Lilly Research Laboratories, Eli Lilly & Company, Lilly Corporate Center, Indianapolis, IN 46285, USA, Guoli Dai, DVM, PhD, Department of Biology, School of Science, Center for Developmental and Regenerative Biology, Indiana University-Purdue University Indianapolis, Indiana 46202. **Grant support:** Translational Research Pilot Grant, Indiana University. **Author contributions:** Conceptualization: BCY, GD, and YW. Investigation: YW and MH. Formal analysis: YM, MH, AC, EW, PK, NC, and SL. Data curation: YW. Writing: YW, AC, and GD. Resources: BCY and GD. **Synopsis** Identifying the primary factors driving liver fibrogenesis is key to the development of effective therapies for chronic liver diseases. We found that hepatic and systematic activin B is rapidly enriched upon the onset of liver injury, and excessive activin B persists as liver injury advances. It directly promotes hepatocyte death and activates hepatic stellate cells, mediating the initiation and progression of hepatic fibrotic response to liver injury. Thus, we for the first time demonstrate activin B as a novel and strong profibrotic factor.

## Abstract

**Background & Aims:** Liver fibrosis is a pivotal pathology in multiple hepatic disease indications, profoundly characterizing disease severity and outcomes. The role of activin B, a TGFβ superfamily cytokine, in liver health and disease is largely unknown. We aimed to investigate whether activin B modulates liver fibrogenesis.

**Methods:** Liver and serum activin B, along with its analog activin A, were analyzed in patients with liver fibrosis from different etiologies and in mouse acute and chronic liver injury models. Activin B, activin A, or both was immunologically neutralized in mice with progressive or established carbon tetrachloride (CCl_4_)-induced liver fibrosis. The direct effects of activin B and A on hepatocytes and hepatic stellate cells (HSCs) were evaluated *in vitro*.

**Results:** As a result, hepatic and circulating activin B was increased in human patients with liver fibrosis caused by several liver diseases. In mice, hepatic and circulating activin B exhibited persistent elevation following the onset of several types of liver injury, whereas activin A displayed transient increases. The results revealed a close correlation of activin B with liver injury regardless of etiology and species. We found that neutralizing activin B largely prevented, as well as remarkably regressed, CCl_4_-induced liver fibrosis, which was augmented by co-neutralizing activin A. Mechanistically, activin B directly promotes hepatocyte death, induces a profibrotic expression profile in HSCs, and stimulates HSCs to form a septa structure. In addition, activin B and A interdependently upregulated the transcription of profibrotic factors including connective tissue growth factor and TGFβ1 in injured livers.

**Conclusions:** We demonstrate that activin B, cooperating with activin A, directly acts on multiple liver cell populations, and drives the initiation and progression of liver fibrosis. Our finding inspires the development of a novel therapy of chronic liver diseases.

## INTRODUCTION

Liver fibrosis is the common consequence of liver injury secondary to a variety of repeated insults and/or injuries including alcoholic liver disease (ALD), non-alcoholic steatohepatitis (NASH), viral hepatitis, and autoimmune liver disease ^1^. The initiation and progression of liver fibrosis is driven by complicated cellular and molecular mechanisms which include a variety of different cell types ^2–5^. Damaged hepatocytes and cytokines released from the resident liver inflammatory cells can directly or indirectly activate hepatic stellate cells (HSCs) into myofibroblasts, leading to the accumulation of collagen I, III and deposition of other extracellular matrix (ECM) components and thus resulting in liver fibrosis ^1, 6, 7^.

Activin proteins consist of dimers formed by four inhibin subunits– inhibin βA, inhibin βB, inhibin βC, and inhibin βE in mammals ^8^. Widely expressed inhibin βA and inhibin βB genes are essential for inducing mesoderm formation during development and for follicle stimulating hormone production ^9–11^. The inhibin βC and inhibin βE are expressed predominantly in the liver, and are dispensable during development and for maintaining adult homeostasis ^12^. Activins A, B, AB, C, and E represent homo or hetero-dimers of inhibin βAβA, βBβB, βAβB, βCβC, and βEβE, respectively ^8^. Activins A, B, and AB signal through activin receptors/Smad2/3 pathway, whereas activins C and E may not ^13^. Activin A is expressed and secreted by hepatocytes and non-parenchymal cells such as HSCs, cholangiocytes and endothelial cells in liver ^14–16^. Several studies demonstrated that activin A induces the activation of HSCs and macrophages and the apoptosis of hepatocytes *in vitro ^14–20^*. An *in vivo* study showed that neutralizing activin A mildly reduced CCl_4_-induced acute liver injury in mice ^21^. However, the association of activins with human liver disease/fibrosis/cirrhosis has yet to be determined.

As structurally related proteins, activin B shares 63% identity and 87% similarity to activin A ^8^. Both ligands bind to the activin receptors II and I, and multiple common AP-1 sites in the promoters of both *inhibin βA* and *inhibin βB* have been identified, which suggests that activin B may act similarly to activin A in mediating liver pathogenesis ^8, 22–24^. Hepatocytes constitutively express abundant inhibin βA but relatively low inhibin βB ^14^. However, hepatic *inhibin βB* gene expression is highly upregulated while *inhibin βA* was downregulated in response to CCl_4_-induced acute liver injury ^25^. Moreover, activin B upregulated hepcidin expression in hepatocytes via Smad1/5/8 signaling in response to several inflammatory insults in mice, whereas activin A did not. These findings suggest a unique role for activin B in mediating liver injury in comparison to activin A ^26^. Taken together, it remains unclear whether activin B is involved in the initiation and progression of liver fibrotic response to liver injury. Thus, the objective of this study was to ascertain the role for activin B to modulate liver fibrogenesis.

## MATERIALS AND METHODS

### Human liver and serum samples

The hepatic mRNA and protein expression of activin A and B was first determined from normal liver (n=5) and the explant which was obtained during liver transplantation from patients with cirrhosis secondary to non-alcoholic steatohepatitis (NASH) (n=8). We further examined the levels of mRNA and protein expression of both activins in patients with different stages (F0-F4) of NASH (n=21). Serum levels of activins were measured in patients with NASH (n=44), excessive drinkers without liver disease (n=36), and those with alcoholic cirrhosis (n=15) compared to normal controls (n=16). All samples were collected under the IUPUI Institutional Review Board-approved protocols.

### Mouse liver injury models

All mouse experiments were performed with the approval of Institutional Animal Care and Use Committee of Eli Lilly and Company and Indiana University-Purdue University Indianapolis. Several mouse models were used for our study. For acute liver injury models, C57BL/6 female mice at the age of 10-12 weeks (Envigo, Indianapolis, IN) received a single intraperitoneal administration of CCl_4_ (Sigma Aldrich, St. Louis, MO) (1:10 dilution in corn oil, 10 ml/kg) for 1.5 hour, 3 hour, 6 hour, 24 hour, and 3 day. For liver fibrosis models, mice were intraperitoneally injected of CCl_4_ twice a week for 4 weeks or 10 weeks ^27, 28^. For ALD model, ethanol oral feeding lasted for 10 days plus binge as described previously ^29^. Bile duct ligation (BDL) model was also used. Briefly, under isoflurane anesthesia (2 to 4 vol %) male C57BL/6 mice (n = 6 to 8) were placed on a heat pad and laparatomized. The common bile duct was exposed, isolated, ligated two times with non-resorbable sutures (polyester 6–0; Catgut, Markneukirchen, Germany). Sham-operated mice underwent a laparotomy with exposure but not ligation of the bile duct. The abdominal muscle and skin layers were stitched, and the mice were treated with ketoprofen as an analgesic. The animals were allowed to recover from anesthesia and surgery under a red warming lamp and were held in single cages. After 14 days mice were euthanized to obtain blood and liver samples.

### Blood biochemistry

Serum aspartate aminotransferase (AST), alanine aminotransferase (ALT), glucose, and total bilirubin levels were measured with a Hitachi Modular Analyzer (Roche Diagnostics, Indianapolis, IN).

### Histology and immunohistochemistry

Formalin-fixed and paraffin-embedded liver sections were subjected to a standard procedure of immunohistochemistry with primary antibodies against F4/80 (eBioscience, San Diego, CA) and myeloperoxidase (MPO, R&D System, Minneapolis, MN). The liver sections were additionally subjected to Masson’s trichrome staining. Images were acquired using digital slide scanning (Aperio Technologies, Vista, CA) and analyzed the stained area percentage by ImageJ (National Institutes of Health, NIH, USA).

### ELISA of activin A and activin B

Activin A and activin B proteins in liver tissue, serum, or cell culture supernatants were quantified by ELISA methods (Activin A ELISA kit, Sigma, St. Louis, MO; Activin B ELISA kit, Ansh labs, Webster, TX) according to the protocols provided by the manufacturers.

### Cell culture

Primary mouse hepatocytes (PMH) were isolated from adult male C57BL/6 mice and were cultured as described (32). LX-2 cells, a human hepatic stellate cell line, was a gift of Dr. Scott L. Friedman from the Mount Sinai School of Medicine (New York, NY). They were cultured in DMEM supplemented with 2% FBS (Gibco, Invitrogen, Carlsbad, CA)

### In situ hybridization

In situ hybridization was performed on formalin-fixed and paraffin-embedded liver sections using mouse inhibin A and inhibin B RNAscope probes and a 2.5 HD Assay-Brown kit (Advanced Cell Diagnostics, Hayward, CA) according to the manufacturer’s protocol (33).

### Microarray Analysis and Quantitative RT-PCR

LX-2 cells were seeded in six well plates with 5 × 10^5^ cells per well and were cultured overnight. The cells were then treated with bovine serum albumin (BSA), activin A, activin B, or TGFβ1 (R&D System, Minneapolis, MN) for six hours, followed by total RNA extraction using TRIzol reagent (Life Technologies, Waltham, MA). Two µg of RNA were reverse transcribed using a cDNA synthesis Kit (Applied Biosystems, Foster City, CA) and were subsequently subjected to HG-U133 plus 2 chips for microarray analysis (Asuragen, Austin, TX). Total RNA was isolated from cultured cells or liver tissues using TRIzol reagents and subsequently was subjected to cDNA synthesis. The qRT-PCR was performed with the cDNAs and the expression of genes of interest was evaluated using TaqMan primer/probe sets with QuanStudio 7 Flex (Applied Biosystems, Carlsbad, CA). GAPDH and RPLPO were used as housekeeping genes.

### Statistical analysis

Statistical significance, *P* < 0.05, was determined by ordinary one-way or two-way ANOVA tests followed by Dunnett’s to compare the differences between experimental and control groups. The data were expressed as means ± S.E.M. GraphPad Prism Software 9.0 was used for data analysis and figure preparation.

## RESULTS

### The levels of hepatic and circulating activin B are significantly increased in patients with liver fibrosis

We first determined whether activin B and A are clinically relevant to different etiologies of liver fibrosis. We found that, in patients with advanced liver fibrosis or cirrhosis, hepatic activin B mRNA and protein exhibited marked increases relative to healthy controls (Figure 1A-B). Circulating activin B did not increase in excessive alcohol users without liver disease but was potently elevated more than fivefold in patients with alcoholic cirrhosis (Figure 1C). In NASH patients, hepatic levels of activin B significantly increased only in those patients with F3 and F4 biopsy confirmed fibrosis compared to F0 controls (Figure 1D) while serum levels of Activin B were increased in those patients with F4 fibrosis (Figure 1E). Additionally, we found that the serum level of activin A markedly increased in those with F1 fibrosis (Figure 1E). Taken together, we demonstrate that the expression of activin B is closely correlated with advanced fibrosis/cirrhosis, irrespective of underlying disease etiologies.

**Figure 1.**
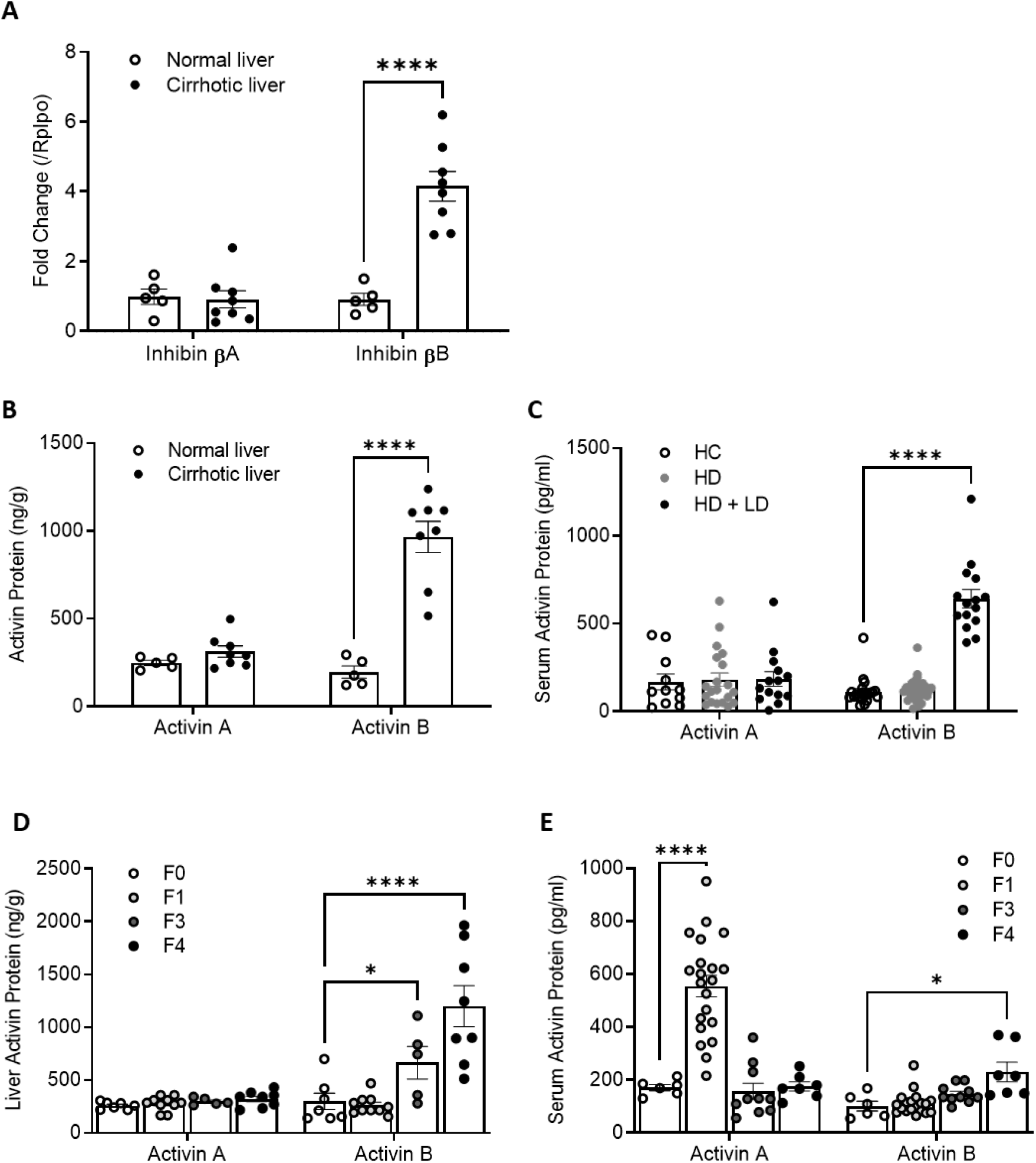
Liver and serum activin B increase in the patients with liver fibrosis. **(A)** The mRNA expression of hepatic inhibin βA and inhibin βB and **(B)** proteins of hepatic activin A and B in patients with cirrhosis (n=8) and healthy controls (n=5) were analyzed by qRT-PCR and ELISA respectively. **(C)** The concentrations of serum activin A and B proteins were determined with ELISA in heathy controls (HC, n=16), heavy drinkers without liver diseases (HD, n=36), and heavy drinkers with liver disease (HD + LD, n=15). Activin A and B proteins were evaluated by ELISA in the **(D)** livers and **(E)** serum of patients with different stages of NASH (F0: n=4, F1: n=6, F3: n=5, and F4: n=6). For all above assays, data are expressed as means ± S.E.M. * *P* < 0.05, **** *P* < 0.0001 compared to heathy controls or F0 group via two-way ANOVA. Note that *inhibin βA* and *βB* encode the subunit of activin A and B respectively.

### The levels of hepatic and circulating activin B are significantly elevated in mouse models of liver injury

To further investigate the expression patterns and cellular sources of activin B and A in liver injury, we performed acute and chronic liver injury studies in mice. In an acute liver injury, mice were administered a single dose of CCl_4_. We found significantly upregulated hepatic *inhibin βB* mRNA expression up to 3 days post injection (Figure 2B), concomitant with the increases in activin B protein concentrations in the sera (Figure 2D) and livers (Figure 2F). In accordance with activin B expression, we observed increases in hepatic mRNA expression, hepatic protein concentration, and serum level of activin A at 24 hours post CCl_4_ injection (Figure 2A, C, and E). To determine these effects following chronic liver injury, we administered mice with CCl_4_ twice weekly for 4 weeks along with a separate ALD model of chronic alcohol plus binge. In the CCl_4_ model, we observed increases in hepatic mRNA expression and serum concentration only for activin B, but not activin A (Figure 3A-B). The cellular sources of activin B and A were revealed via in situ hybridization. Activin B and A were mainly transcribed in hepatocytes and biliary epithelial cells in vehicle-controlled livers and additionally in fibrogenic cells in the fibrotic livers (Figure 3C). Similar findings were found in mice fed with chronic alcohol plus binge (Figure 3D-E). Activin B expression was quantified in bile duct ligation (BDL) model to further characterize activin B response to liver damage. As quickly as 6 hours post-surgery, mice had elevated circulating activin B protein, and this response persisted up to 7 days after BDL (Figure 4A). Similarly, hepatic *inhibin βB* exhibited persistent upregulation during the first 7 days after surgery (Figure 4B). In contrast, during the same period, circulating activin A protein levels were not significantly altered (Figure 3A) and hepatic *inhibin βA* mRNA expression showed a transient increase at 3 days following surgery (Figure 3B). Collectively, we demonstrate that, irrespective of liver injury types, activin B is persistently associated with liver disease progression from acute phase to chronic phase, whereas activin A is transiently relevant to the acute phase. Moreover, the association of activin B with liver fibrosis is highly conserved between humans and mice.

**Figure 2.**
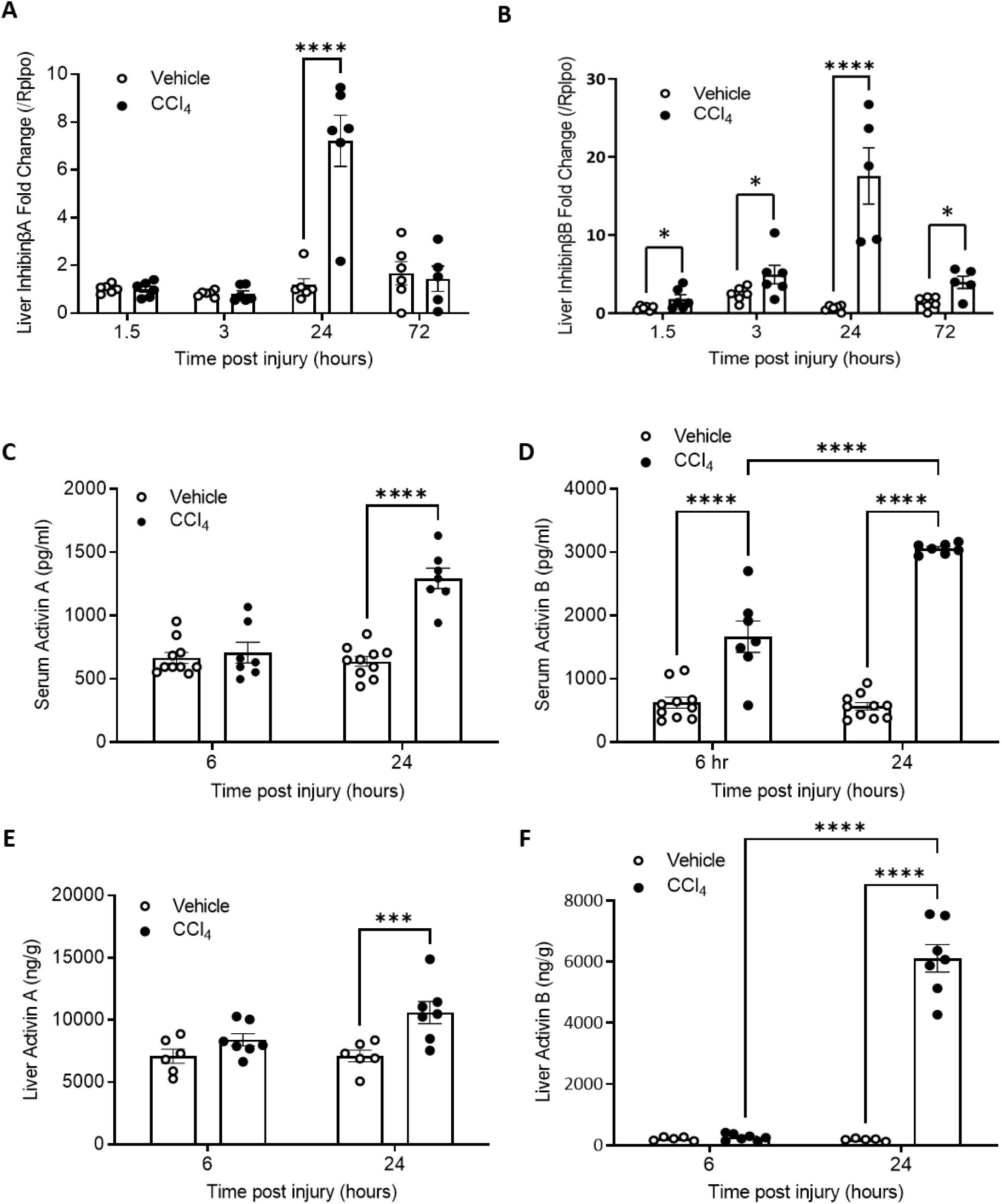
Liver and serum activin B increases in mice following acute liver injury. Mice were given a single administration of CCl_4_ and tissues were collected at the indicted times post injury. The mRNA expression of liver **(A)** *inhibin βA* and **(B)** *inhibin βB* was analyzed by qRT-PCR at indicated time points (n=8). **(C&D)** Serum and **(E&F)** liver activin proteins were quantified via ELISA at the indicated time points (n=8). For all above quantitative assays, data are expressed as means ± S.E.M. *** *P* < 0.001, ***** *P* < 0.0001 via two-way ANOVA relative to vehicle controls.

**Figure 3:**
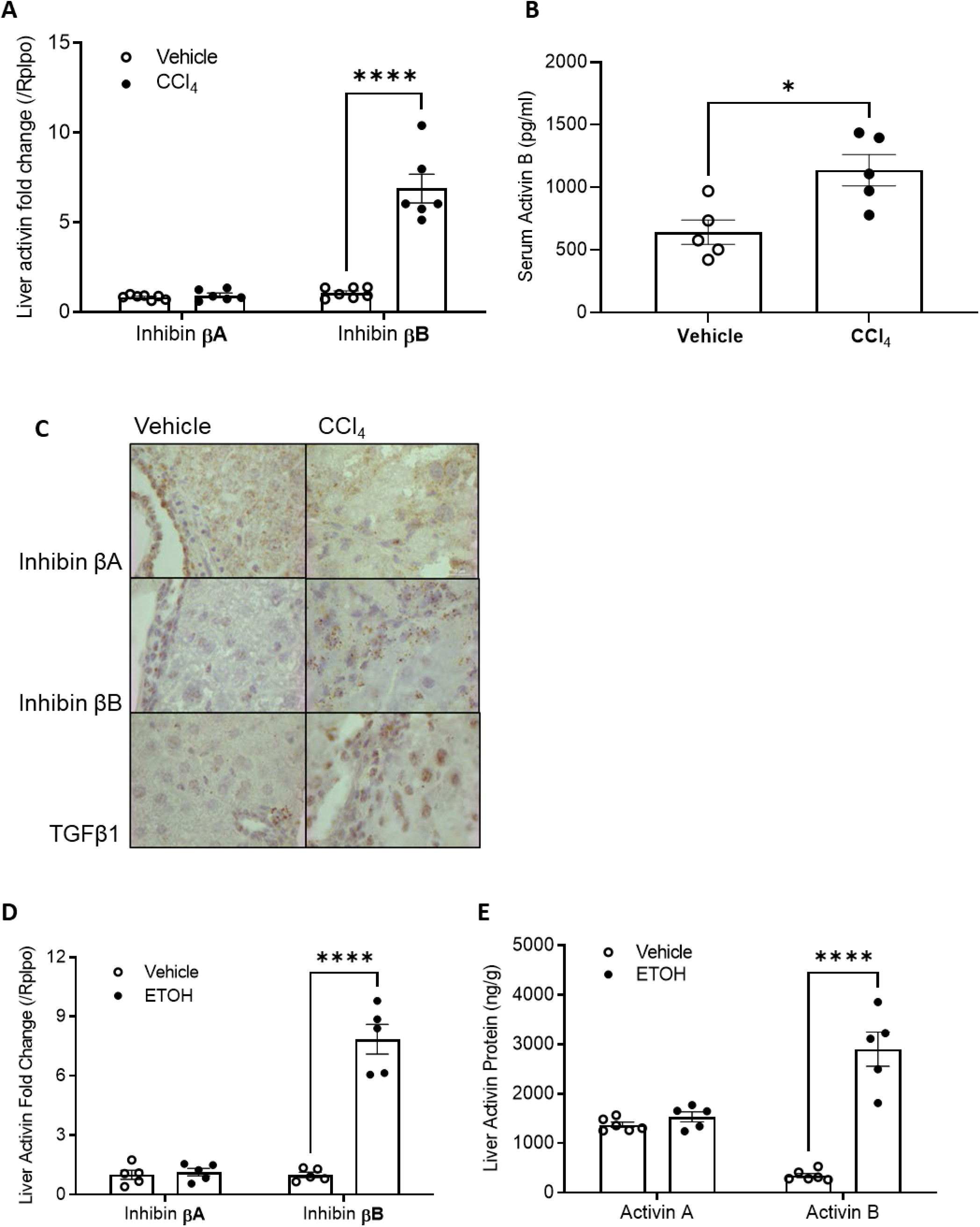
mRNA and protein levels of activin B and A in CCl_4_–induced chronic liver injury. Carbon tetrachloride (CCl_4_) or vehicle was dosed twice per week for 4 weeks in mice, **(A)** mRNA expression of hepatic *inhibin βA* and *inhibin βB* was assessed by qRT-PCR (n=10); **(B)** concentrations of serum activin B protein were quantified via ELISA (n=10); and **(C)** inhibin βA-, inhibin βB-, and TGFβ1-expressing cells were visualized with in situ hybridization on liver sections using mouse inhibin A and inhibin B RNAscope probes and a 2.5 HD Assay-Brown kit. In a separate study, following 10 days post-oral alcohol (ETOH) plus binge administration in mice, **(D)** hepatic *inhibin βA* and *inhibin βB* transcript levels were determined by qRT-PCR (n=7), and **(E)** hepatic activin A and activin B protein contents were quantified via ELISA (n=7). For all above quantitative assays, data are expressed as means ± S.E.M. * *P* < 0.05, **** *P* < 0.0001 via two-way ANOVA relative to vehicle controls.

**Figure 4:**
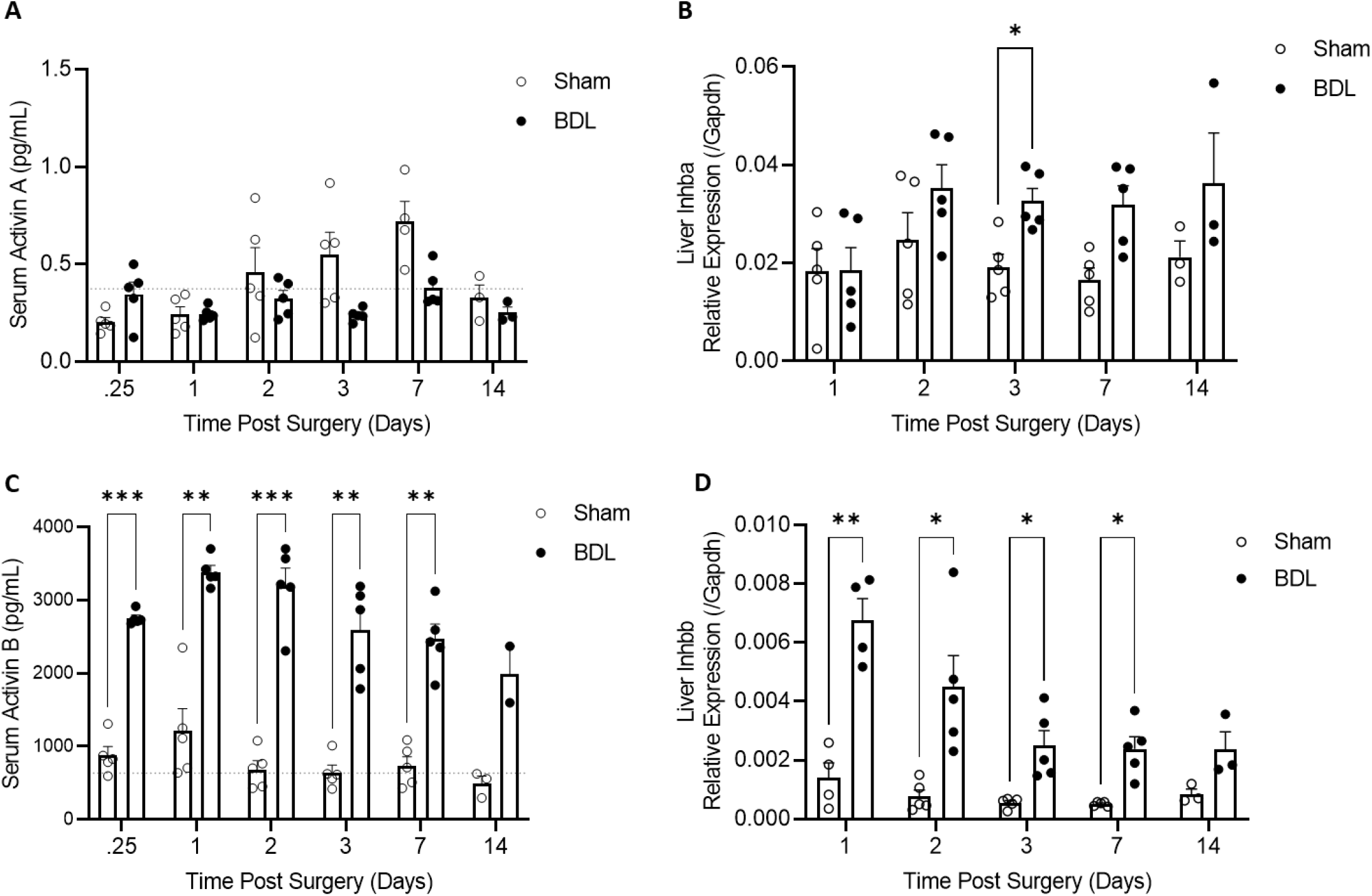
mRNA and protein levels of activin B and A in bile duct ligation (BDL)-induced chronic liver injury. Adult male C57bl/6 mice underwent BDL surgery with age matched sham laparotomy surgery controls. Animals were sacrificed 0.25, 1, 2, 3, 7, and 14 days post-surgery. **(A)** Serum activin A and **(C)** activin B protein contents were quantified via ELISA along with liver mRNA expression of **(B)** *inhibin βA* and **(D)** *inhibin βB* via qRT-PCR (n=5 for all time points except day 14; n=3 for day 14). Age matched naïve control animals were sacrificed on day 14 and mean serum activin protein levels are expressed as grey dotted lines (activin A 0.3745 pg/mL ± 0.1364; activin B 636.3 pg/mL ± 66.11; n=4). All other data are expressed as means ± S.E.M. * *P* < 0.05, ** *P* < 0.01, *** *P* < 0.001 via two-way ANOVA relative to sham controls collected at the same time point.

### Neutralization of activin B prevents CCl_4_-induced liver fibrosis

The above association studies in humans and mice strongly suggested that activin B and A participate in mediating liver fibrogenic response to liver injury. Therefore, we set out to test inhibition of each individual ligand pharmacologically to evaluate their separate and combined contributions to hepatic fibrogenesis. Global gene knockouts of these two widely produced activin ligands cause developmental defects, reproductive failure, or postnatal death in mice (^9–11^). We thereby used neutralizing antibodies to systemically inactivate these two proteins and subsequently examine their effects on the initiation of CCl_4_-induced liver fibrosis. There were five treatment groups: (1) vehicle; (2) IgG + CCl_4_; (3) activin A antibody + CCl_4_; (4) activin B antibody + CCl_4_; and (5) combination of both antibodies + CCl_4_. In the initial association studies, we found time windows during which both activin A and B were induced in the acute phase of liver injuries (Figure 2A-F). This co-induction suggested a possible spatiotemporal coordination between the two activin ligands, warranting combination antibody treatment in this study. Antibodies were initially dosed half an hour before the first CCl_4_ injection and were dosed weekly thereafter. A dosage of 10 mg/kg of activin A antibody weekly was used because our previous study demonstrated the greatest efficacy of this regimen in regressing degeneration of injured skeletal muscle in mice ^30^.

As a result, Activin B antibody exerted extensive beneficial effects, including reduced liver injury indicated by serum ALT and AST (Figure 5A-B), improved liver functions estimated by serum glucose and total bilirubin levels (Figure 5C-D), and decreased liver fibrosis analyzed by collagen staining and collagen 1α1 mRNA expression (Figure 5E-G). Activin A antibody treatment reduced liver injury and improved liver functions to a lesser extent compared with activin B antibody, but did not decrease total bilirubin and liver fibrosis, although collagen 1α1 mRNA expression was inhibited (Figure 5A-G). The dual antibodies showed beneficial effects equivalent to, or in some cases greater than, activin B antibody alone (Figure 5A-G). In CCl_4_ chronically damaged livers, activin B and A are essential collaborators to induce profibrotic genes chemokine (C-X-C motif) ligand 1 (CXCL1), cytokine-inducible nitric oxide synthase (iNOS), connective tissue growth factor (CTGF), and TGFβ1, because neutralizing either one of them prevented or inhibited the upregulation of these genes (Figure 5H). Together, these data demonstrate that (1) activin B and, to a much lesser extent, activin A mediate the initiation of liver fibrosis and (2) activin B inhibition or, even better, both activin B and A inhibition largely prevents liver fibrosis.

**Figure 5:**
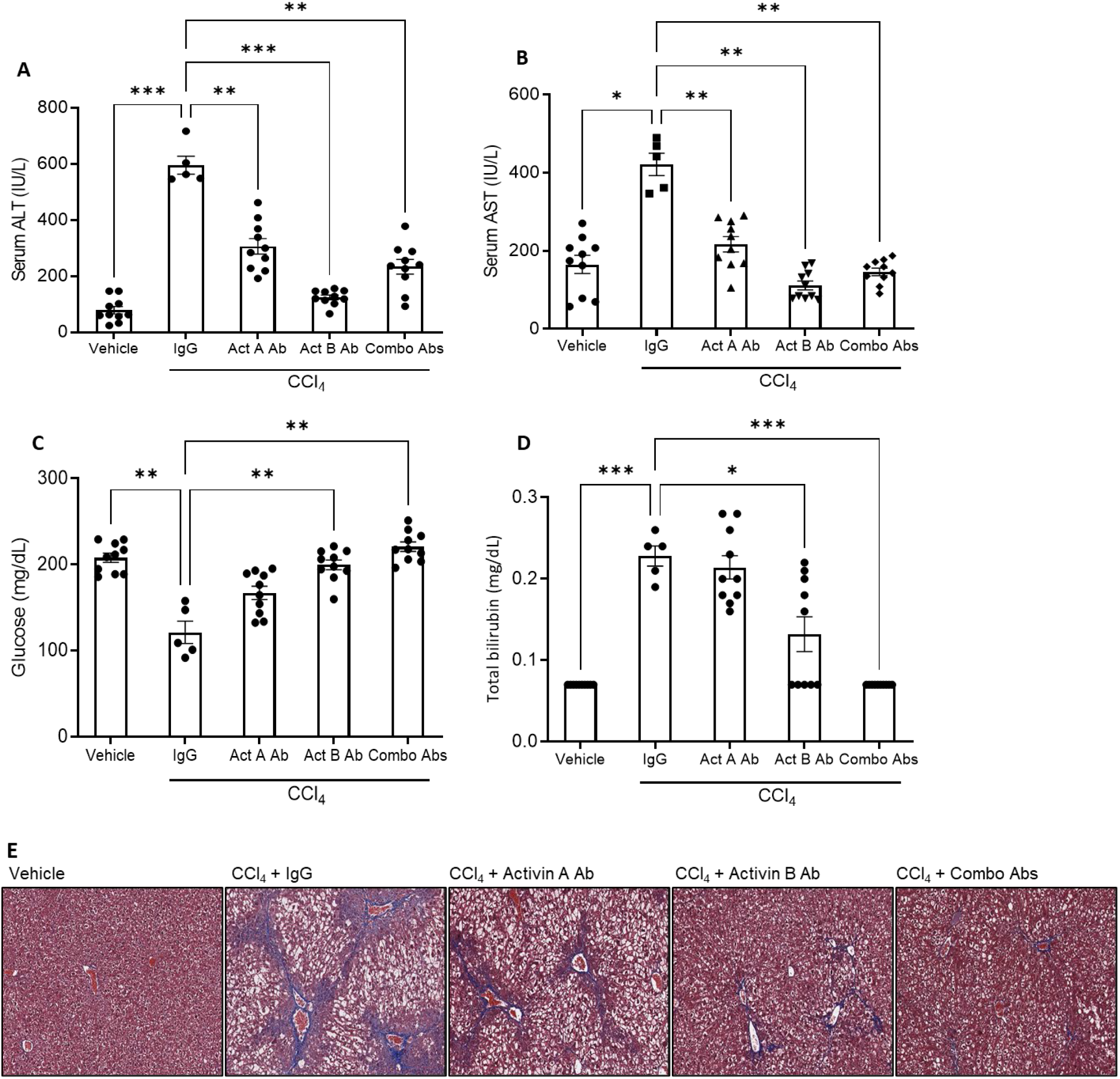
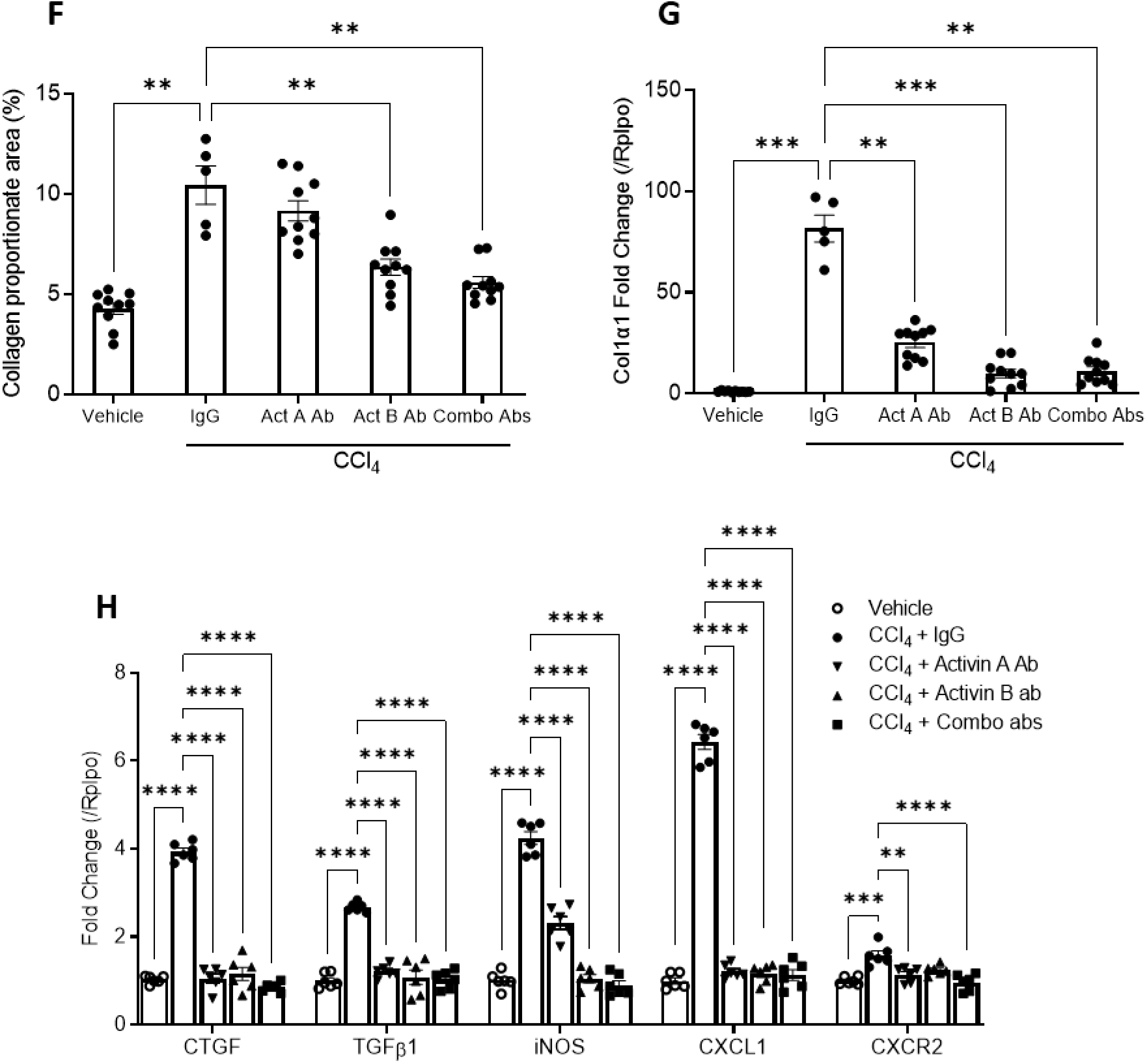
Treatments with activin B antibody, activin A antibody, or combination of them display distinct effects in preventing liver fibrosis induced by CCl_4_ in mice. Adult female mice were administered carbon tetrachloride (CCl_4_) or corn oil vehicle twice per week for 4 weeks. Half an hour before the first CCl_4_ injection each week, mice were treated with IgG (60 mg/kg), activin A antibody (10 mg/kg of activin A antibody + 50 mg/kg of IgG), activin B antibody (50 mg/kg of activin B antibody + 10 mg/kg of IgG), or combination of activin A and activin B antibodies (10 mg/kg of activin A antibody + 50 mg/kg of activin B antibody). Four weeks after the initial CCl_4_ injection, **(A)** ALT, **(B)** AST, **(C)** glucose, and **(D)** total bilirubin were analyzed in blood. **(E)** Representative liver sections stained with **(E)** Masson’s trichrome. **(F)** Percent Masson’s trichrome staining areas were quantified by ImagJ. **(G)** The mRNA expression of hepatic Col1α1 was evaluated by qRT-PCR. **(H)** Transcripts of the genes indicated were quantified by qRT-PCR in liver tissues. Data are expressed as means ± S.E.M. (n=10). * *P* < 0.05, ** *P* < 0.01, *** *P* < 0.001, and **** *P* < 0.0001 compared to IgG controls via ordinary one-way ANOVA for figures A-D & and F-G. Statistical analysis was performed via two-way ANOVA following by Dunnett’s multiple comparisons test relative to IgG controls for panel H. Total bilirubin levels below the limit of detection (0.14 mg/dL) were replaced by half of the limit of detection (0.07 mg/dL) for statistical analysis.

### Neutralization of activin B regresses established CCl_4_-induced liver fibrosis

The superior effects of antibody-mediated inactivation of activin B or both activin B and A in preventing liver fibrosis prompted us to test the same strategy to reverse existing liver fibrosis. Following the same study design as the prevention study, CCl_4_ was injected twice per week for ten continuous weeks. Starting at the seventh week when liver fibrosis was fully established, antibodies were dosed weekly for the remaining four weeks. Consequently, we found distinct reversal effects among the antibody treatment groups. The reversal effects followed the sequence of magnitude of effect where inactivating both activin B and A had greater effect than inactivating activin B alone, and inactivating activin A had the lowest effect. The combinational inactivation exerted the most beneficial effects across all assessments, including reduced liver injury assessed by serum ALT and AST (Figure 6A-B), improved liver function estimated by serum glucose and total bilirubin levels (Figure 6C-D), decreased collagen deposition (Figure 6E-F), and less macrophage infiltration (Figure 6G and 6I). Inactivating activin B alone generated a stronger anti-fibrotic effect (Figure 6E-F), but nearly equal effects in other assessments (Figure 6A-D, G-I), compared with inactivating activin A alone. Of note, inactivating activin B alone and inactivating both activin B and A equivalently regressed liver fibrosis (Figure 6E-F). Neutrophils (MPO-positive cells) and Kupffer cells (F4/8-positive cells) were similarly distributed in fibrotic livers, concentrating in septa. Neutralizing activin A, activin B, or both did not alter the total number of infiltrated neutrophils, but almost equally reduced the total number of Kupffer cells (Figure 6G-I). Collectively, these results demonstrate that (1) activin B is a stronger driver of liver fibrogenesis than activin A; (2) these two activin ligands may act cooperatively to promote the progression of chronic liver injury; and (3) neutralization of activin B, or both of these ligands, largely reverses liver fibrosis.

**Figure 6:**
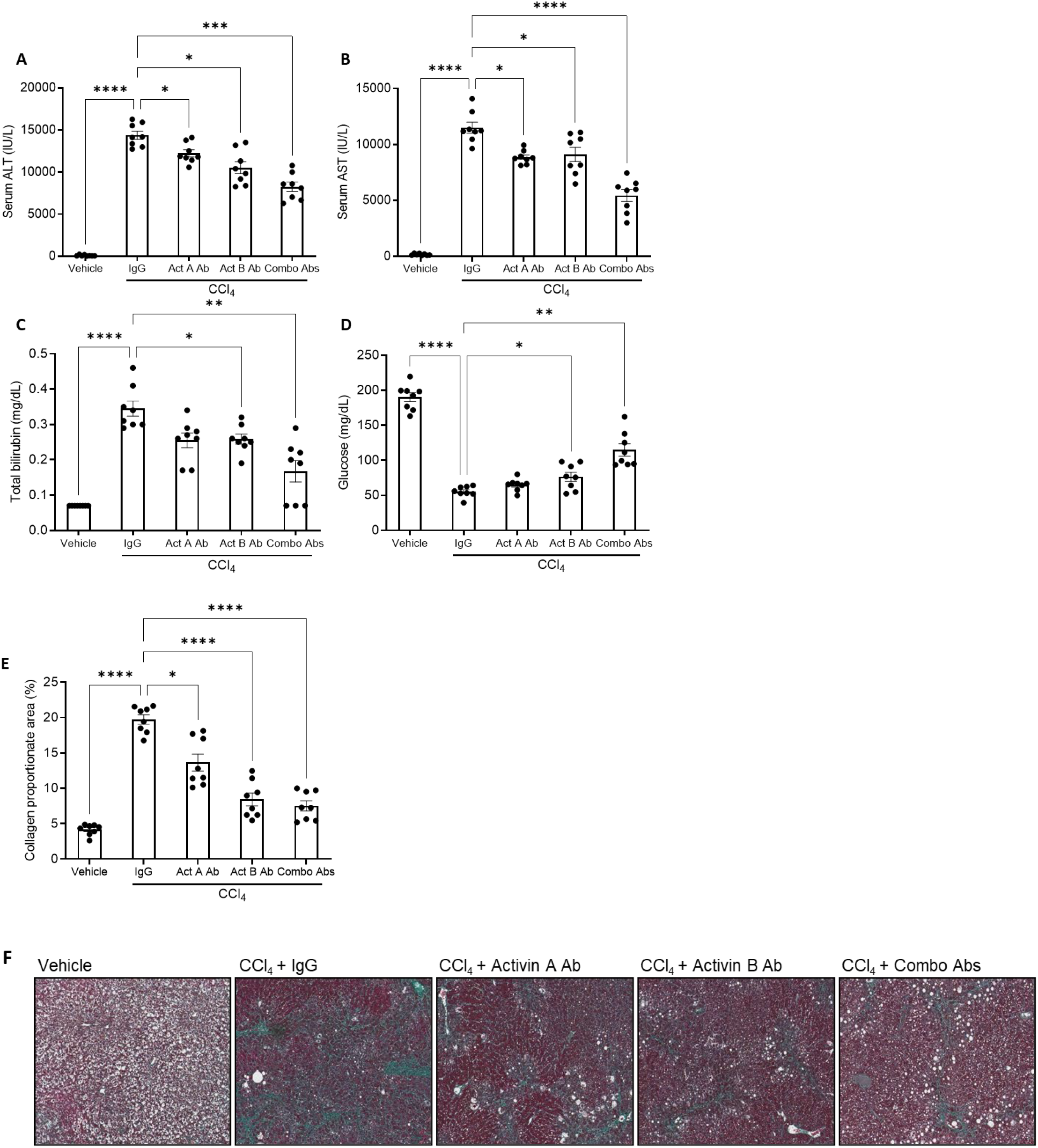
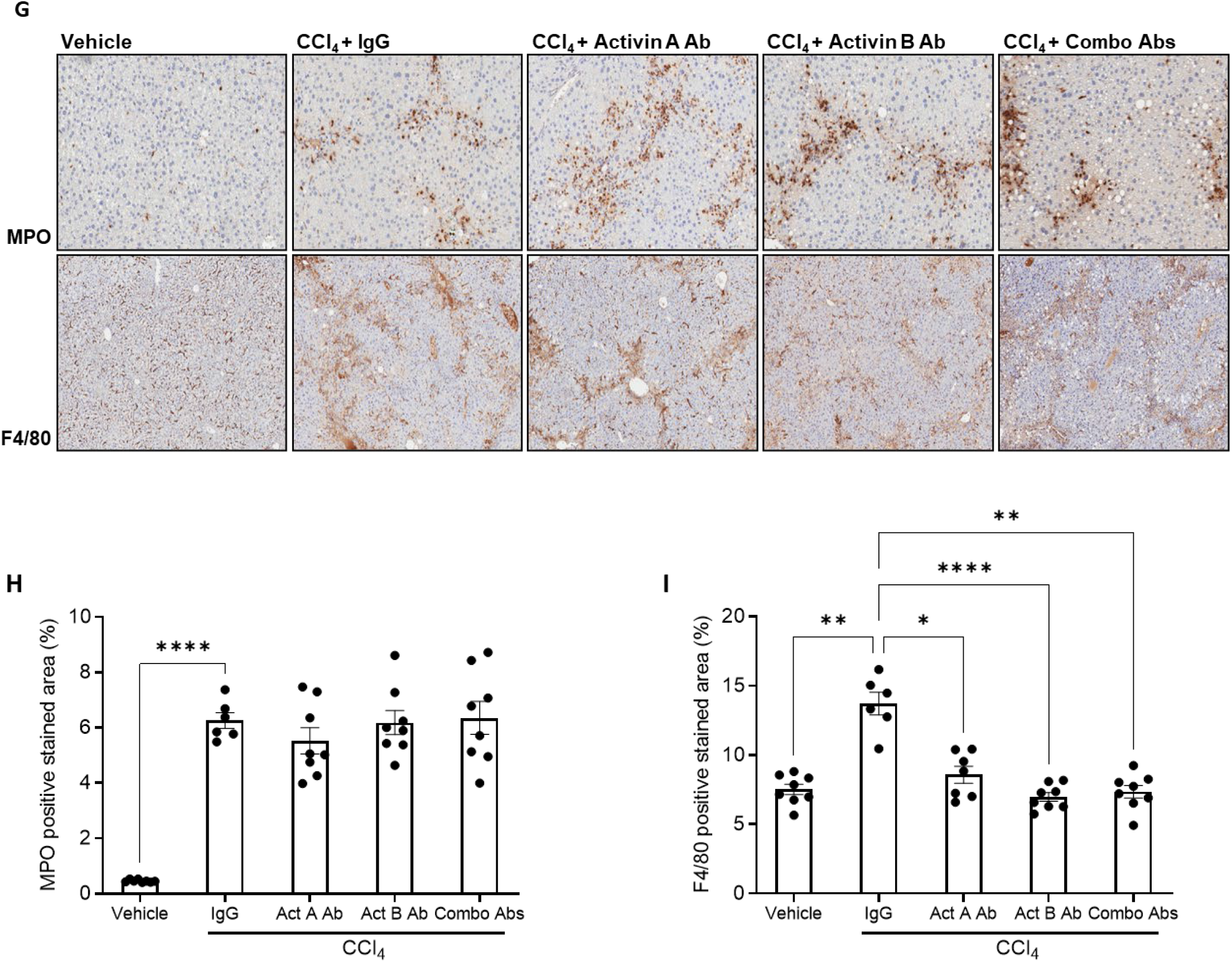
Activin B antibody, activin A antibody, and combination of them display different effects in regressing liver fibrosis induced by CCl_4_ in mice. Adult female mice were subjected to carbon tetrachloride (CCl_4_) or vehicle injection twice per week for 10 weeks. Starting from the seventh week, these mice were treated with IgG (60 mg/kg), activin A antibody (10 mg/kg of activin A antibody + 50 mg/kg of IgG), activin B antibody (50 mg/kg of activin B antibody + 10 mg/kg of IgG), or combination of activin A and activin B antibodies (10 mg/kg of activin A antibody + 50 mg/kg of activin B antibody) weekly. Ten weeks after the initial CCl_4_ injection, **(A)** ALT, **(B)** AST, **(C)** glucose, and **(D)** total bilirubin were analyzed in blood **(E)** Masson’s trichrome collagen staining areas were quantified by ImagJ. <u>**(F)** Representative</u> liver sections stained with Masson’s trichrome. **(G)** Representative liver sections immunostained with MPO or F4/80 antibody. **(H&I)** quantification of percent positive staining area of **(H)** MPO and **(I)** F4/80. Data are presented as means ± S.E.M. (n=8). * *P* < 0.05, ** *P* < 0.01, *** *P* < 0.001, and **** *P* < 0.0001 compared to IgG controls via ordinary one-way ANOVA relative to IgG controls. Total bilirubin levels below the limit of detection (0.14 mg/dL) were replaced by half of the limit of detection (0.07 mg/dL) for statistical analysis.

### Activin B is produced by injured primary hepatocytes and in turn promotes their death

*In situ* hybridization showed that hepatocytes abundantly expressed *inhibin βB* mRNA in injured liver (Figure 3C). To examine whether activin B and A proteins are secreted by hepatocytes and how these proteins respond to hepatocyte injury, we exposed primary mouse hepatocytes (PMH) to CCl_4_ or lipopolysaccharide (LPS). We found that CCl_4_ damaged these cells and induced their necrosis, which was indicated by reduced cell viability and increased release of ALT and AST into the culture supernatants (Figure 7A-B). Injured hepatocytes increased the secretion of both activin B and A proteins (Figure 7C). Cell viability was significantly, further significantly, and additively improved by neutralizing activin A, activin B, and both of them, respectively (Figure 7D). In contrast, LPS only stimulated PMH to increase activin A production without affecting activin B, ALT and AST (Figure 7B-C). These data suggest that hepatocytes increase their production of activin B and A in a toxin-dependent manner. Of note, these two activins augmented hepatocyte injury, as neutralization of these proteins ameliorated cell viability following insults (Figure 7D). We also found that PMH responded to individual or combinational treatment of these two proteins by equivalently upregulating the transcription of genes encoding profibrotic factors TGFβ1, CTGF, and Col1α1, as well as by distinctly altering the mRNA expression of skeletal muscle alpha actin *ACTA1*, *Smad3*, and *IKBKB* (Figure 7E). Thus, the results suggest that activin B and A have redundant, specific, and interactive actions on hepatocytes. Taken together, our data suggest that injured hepatocytes produce activin B and A, which in turn promote hepatocyte death and fibrotic response.

**Figure 7:**
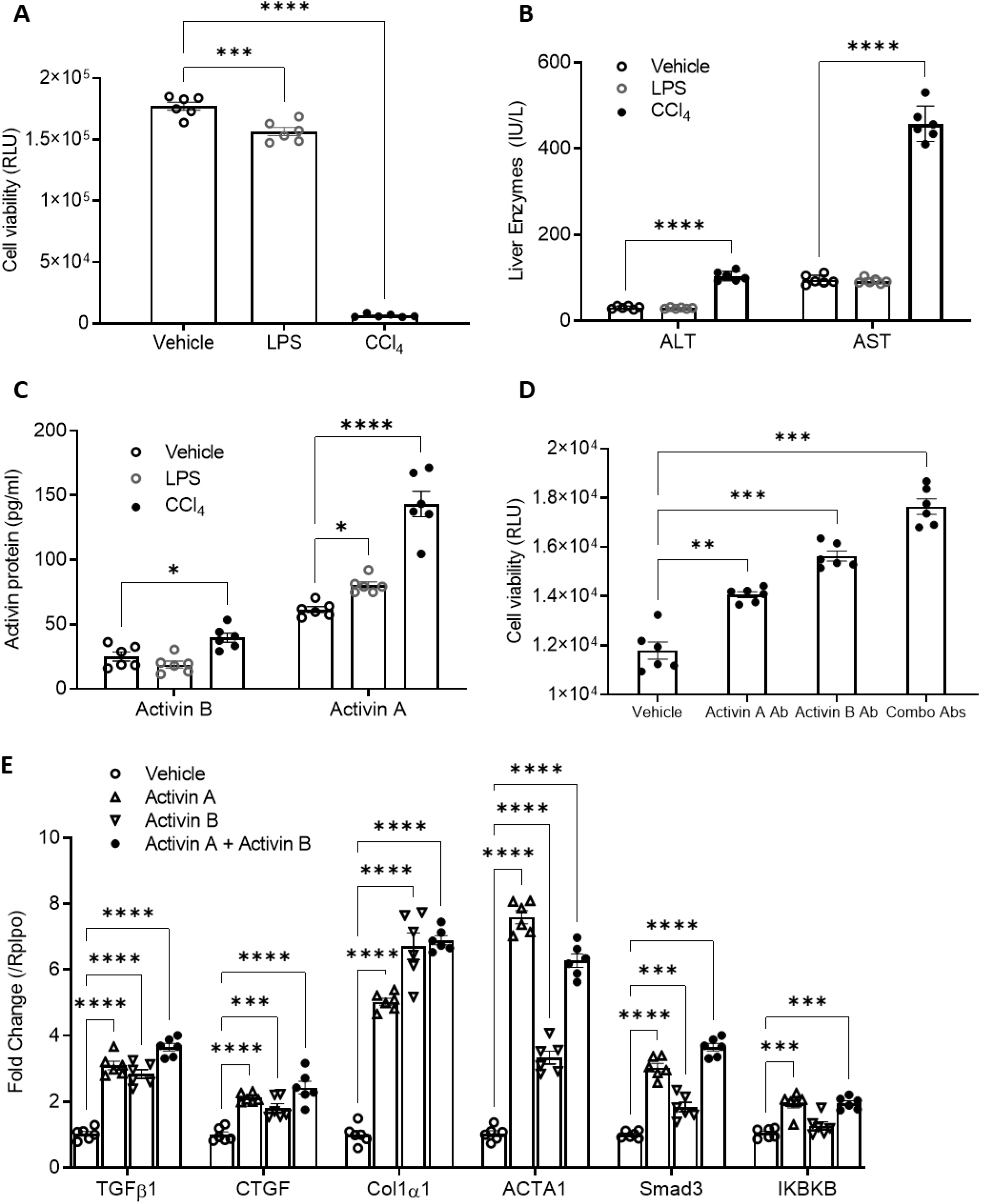
Activin B is produced by injured primary mouse hepatocytes and induces expression of genes encoding fibrotic factors in these cells. Primary mouse hepatocytes (PMHs) were isolated from adult male mice and cultured overnight. Subsequently, the cells were treated with vehicle (corn oil), lipopolysaccharide (LPS, 10 µg/ml), or 0.5% CCl_4_ for 24 hours. **(A)** Cell viability, **(B)** supernatant ALT and AST, and **(C)** supernatant activin A and B proteins were analyzed. **(D)** Cell viability of primary hepatocytes following 24 hours of treatment with 0.5% CCl_4_ and co-treatment with IgG, activin A antibody, activin B antibody, or combination of both antibodies (100 ng/ml each). **(E)** mRNA expression of various genes was evaluated by qRT-PCR in PMHs treated with activin A or activin B (100 ng/ml each), or their combination for 24 hours. For all above assays, data are expressed as means ± S.E.M. * *P* < 0.05, ** *P* < 0.01, *** *P* < 0.001, and **** *P* < 0.0001 via Ordinary one-way ANOVA for panels A & D and two-way ANOVA for panels B, C, & E compared to vehicle controls.

### Activin B directly promotes HSC activation

Myofibroblasts centrally drive liver fibrogenesis and are primarily differentiated from activated HSCs ^31^. LX-2 cells, a human HSC cell line, largely mimic the behavior of primary HSCs and thereby have been widely used to study the property of HSCs ^32^. Because LX-2 cells are more relevant to humans than mouse HSCs, we tested the behavioral responses of LX-2 cells to activin A, activin B, combination of both, and TGFβ1, a recognized regulator of HSC activity. We found that LX-2 cells formed a septa-like structure following 24 hours exposure to these three ligands (Figure 8A), mimicking the behavior of HSCs during liver fibrogenesis. This observation demonstrates that activin B and A directly activate HSCs. This finding also urged us to examine whether these activin ligands redundantly act on HSCs at a molecular level. Hence, we treated LX-2 cells with activin A, activin B, or TGFβ1 protein for six hours, and profiled their early responsive genes by microarray analysis. As a result, these three proteins regulate overlapping but differential gene networks in these cells (Figure 8B). The overlapping 877 genes were associated predominately with HSC activation and hepatic fibrosis, including upregulated TGFβ signaling negative feedback modulator transmembrane prostate androgen-induced protein (TMEPAI), early growth response protein 2 (EGR2), calcium ion-binding protein matrix gla protein (MGP), and downregulated BMP4, dual specificity phosphatase 6 (DUSP6), extracellular matrix glycoprotein TNXB, IL-8, and IL-17 receptor C (Figure 8C-D). These data suggest that activin signaling dictates a spectrum of HSC properties redundantly via multiple ligands including activin A and B. On the other hand, each of these individual ligands has a large and unique set of target genes associated with critical cellular functions. These data together suggest that activin B is a novel direct regulator of HSCs and that activin ligands distinctly but coordinately modulate the transcriptome of HSCs.

**Figure 8:**
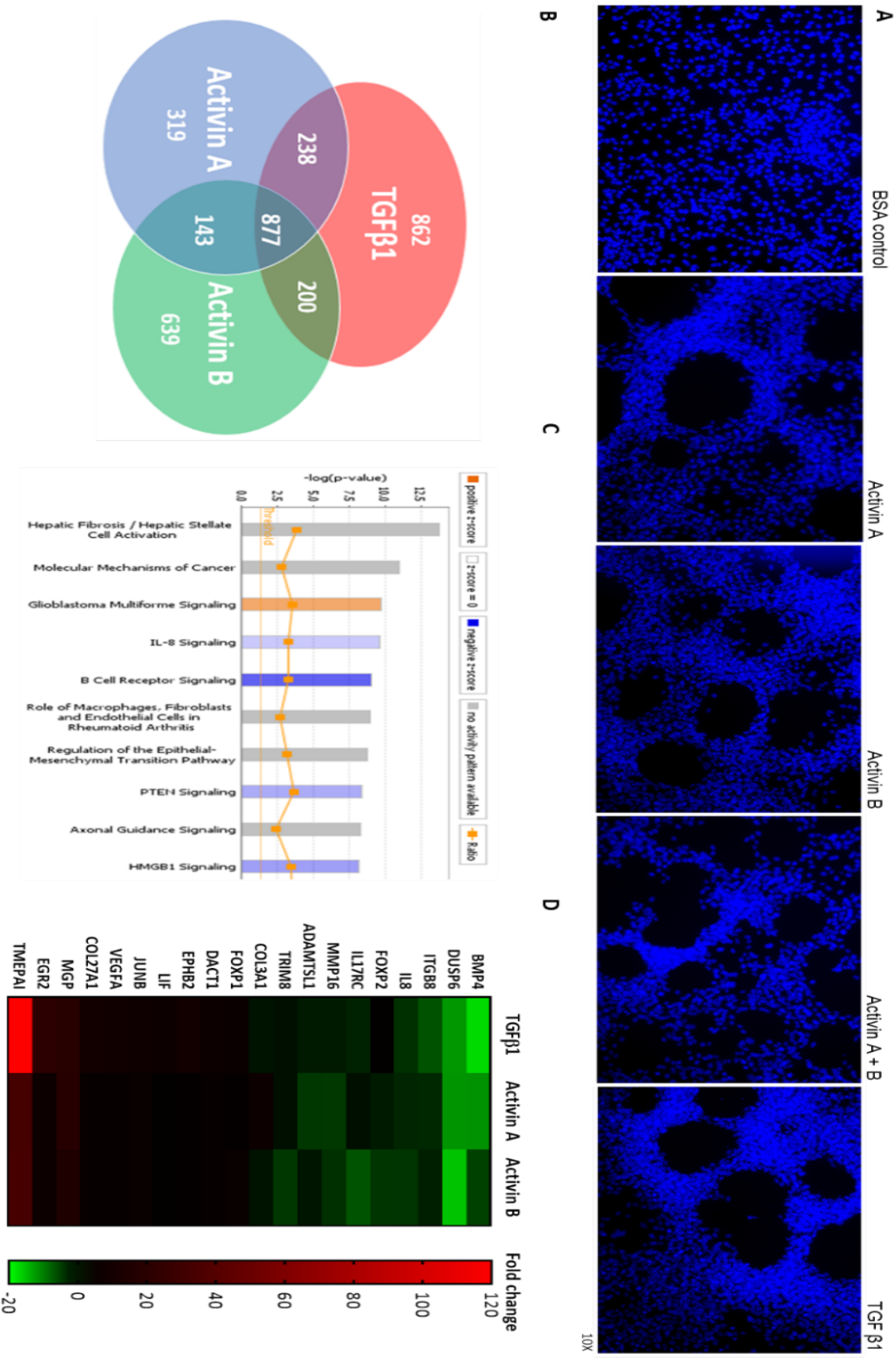
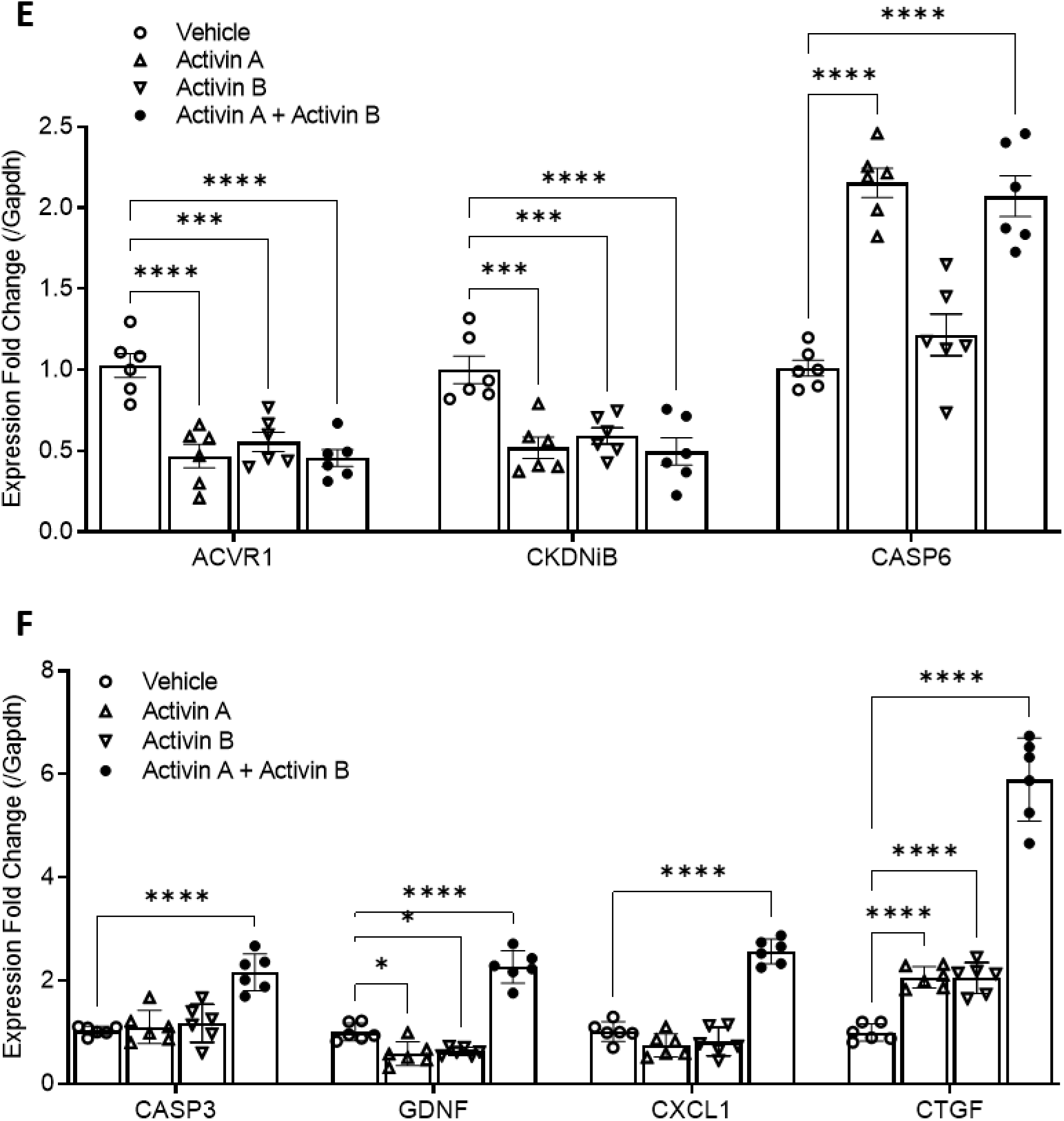
Activin B morphologically and molecularly activates HSCs. **(A)** LX-2 cells were treated with bovine serum albumin (BSA, 100 ng/ml), activin A (100 ng/ml), activin B (100 ng/ml), their combination (100 ng/ml each), or TGFβ1 (5 ng/ml) for 24 hours and then underwent 4′, 6-diamidino-2-phenylindole (DAPI) staining. **(B)** LX-2 cells were treated with activin A (100 ng/ml), activin B (100 ng/ml), or TGFβ1 (5 ng/ml) for 6 hours. Total RNAs were isolated, reverse transcribed to cDNA, and then subjected to microarray analysis using HG-U133 plus 2 chips (n=6). Pie chart shows the numbers of genes commonly or uniquely regulated by the individual ligands. **(C)** The top ten signaling pathways revealed by Ingenuity Canonical Pathway analysis of the 877 target genes shared by these three ligands. **(D)** Heat-map of the 20 genes exhibiting the highest magnitudes of upregulation or downregulation in response to these three ligands. **(E&F)** LX-2 cells were treated with vehicle, activin A (100 ng/ml), activin B (100 ng/ml), or their combination (100 ng/ml of each) for 24 hours. The expression of the genes indicated was assessed with qRT-PCR. Data are shown as means of fold changes relative to vehicle controls ± S.E.M. * *P* < 0.05, ** *P* < 0.01, *** *P* < 0.001, and **** *P* < 0.0001 via two-way ANOVA compared to vehicle controls

To gain insight into how activin A and B interactively act on HSCs, we exposed LX-2 cells to activin A or B alone or both and subsequently examined how a group of genes known to regulate HSC activity respond transcriptionally. We observed four scenarios: (1) *ACVR1* (activin A receptor type 1) and *CKDN1B* (cyclin-dependent kinase inhibitor 1B) equivalently responded to individual ligand (Figure 8E); (2) *CASP6* (caspase-6) solely responded to activin A (Figure 8E); (3) *CASP3*, *GDNF* (glial cell line-derived neurotrophic factor), and *CXCL1* specifically responded to dual ligands (Figure 8F); and (4) *CTGF* equally responded to individual ligands but synergistically to dual ligands (Figure 8F). The results indicate that activin B and A have redundant, unique, and interactive effects on HSCs. Taken together, these *in vitro* data demonstrate that activin B and A directly, redundantly, as well as cooperatively promote the activation of HSCs.

## DISCUSSION

Our studies revealed that circulating activin B is closely associated with liver injury and fibrosis regardless of etiologies and species. These data suggest a highly conserved, activin B-mediated mechanism of response to liver insults in mammals. This response is activated rapidly following liver injury, operates stably as the progression of disease, and predominates throughout liver fibrosis. Increased mRNA and protein levels of hepatic activin B is always concomitant with enriched circulating activin B in patients and mice with liver injury. Therefore, it is likely that increased production of hepatic activin B largely contributes to its systemic elevation during liver fibrosis development. This warrants further investigation to potentially develop Activin B as a reliable and sensitive serum marker for monitoring liver injury progression.

We demonstrated that activin B acts as a potent driver of the complications (hepatocyte injury and fibrosis) of chronic liver injury through *in vitro* and *in vivo* studies. Damaged hepatocytes produced activin B that facilitated hepatocyte death in an autocrine manner. This *in vitro* finding may explain why activin B *in vivo* inactivation reduced the release of ALT and AST from hepatocytes in injured liver. We also demonstrated that activin B acts as a strong pro-fibrotic factor following liver injury. *In vitro*, activin B altered the transcriptome of HSCs towards a pro-fibrotic myofibrolast-like profile. These effects were highlighted by upregulated expression of genes encoding profibrotic factors including collagens, matrix metalloproteinase, and CTGF. Activin B also induced HSCs to form a septa-like structure *in vitro*. We for the first time revealed the property of these cells, offering an *in vitro* assay to investigate the behavior and morphology of these cells in fibrotic liver. *In vivo*, neutralizing activin B alone largely repressed septa formation, collagen deposition, and expression of fibrotic genes such as CTGF and TGFβ1 in chronically injured liver. Microarray data have provided us with a list of activin B target genes of interest for further investigation to elucidate how activin B modulates HSC activity during liver injury. Collectively, these findings allow us to propose that activin B promotes the initiation and progression of liver fibrosis by augmenting hepatocyte death and sustaining HSC activation.

Our studies discovered that increased activin B requires the presence of activin A to optimally mediate liver injury progression. We showed that hepatic and circulating activin A were only transiently increased during the acute phase of liver injury and were kept at pre-injury levels throughout the long-term chronic phase. However, neutralizing activin A alone did produce some beneficial effects, although significantly less than neutralizing activin B alone, in both prevention and reversal studies. In addition, both activin B and A were required for the upregulation of profibrotic factors CTGF and TGFβ1 in chronically damaged livers, because neutralizing either one of the two TGFβ ligands fully abolished the effects. Moreover, we observed several additive or interdependent effects between activin B and activin A *in vitro* on HSCs as well as *in vivo* in fibrotic livers. Most notably, inactivating both activin B and activin A gave rise to the most profound beneficial effects across hepatic structural and functional assessments compared to inactivating activin B or A alone. These observations enable us to reason that, as liver injury progresses, elevated activin B needs constitutive activin A for cooperative pathological actions, since both ligands are required for activating certain cellular programs that otherwise would not be initiated by a single ligand. This represents a novel mode of action of activin ligands in general and a new mechanism governing the actions of activin B and A specifically. This discovery enables us to gain important mechanistic insight into the actions of activin B and A during fibrogenesis. Our pre-clinical studies demonstrate that targeting activin B or ideally both activin B and A is a promising strategy to prevent and even reverse liver fibrosis. This finding warrants future investigations to evaluate the efficacy of Activin B blockade in diverse liver injury models.

In summary, we identified activin B as a critical promoter and a promising therapeutic target of liver fibrosis. The results of our studies permits us to propose a working hypothesis depicted in Figure 9. This hypothesis highlights activin B as a key profirotic factor driving liver fibrogenesis and its advance and lays the groundwork for future investigations to elucidate how activin B acts in various liver cell populations during the initiation and progression of liver fibrosis.

**Figure 9:**
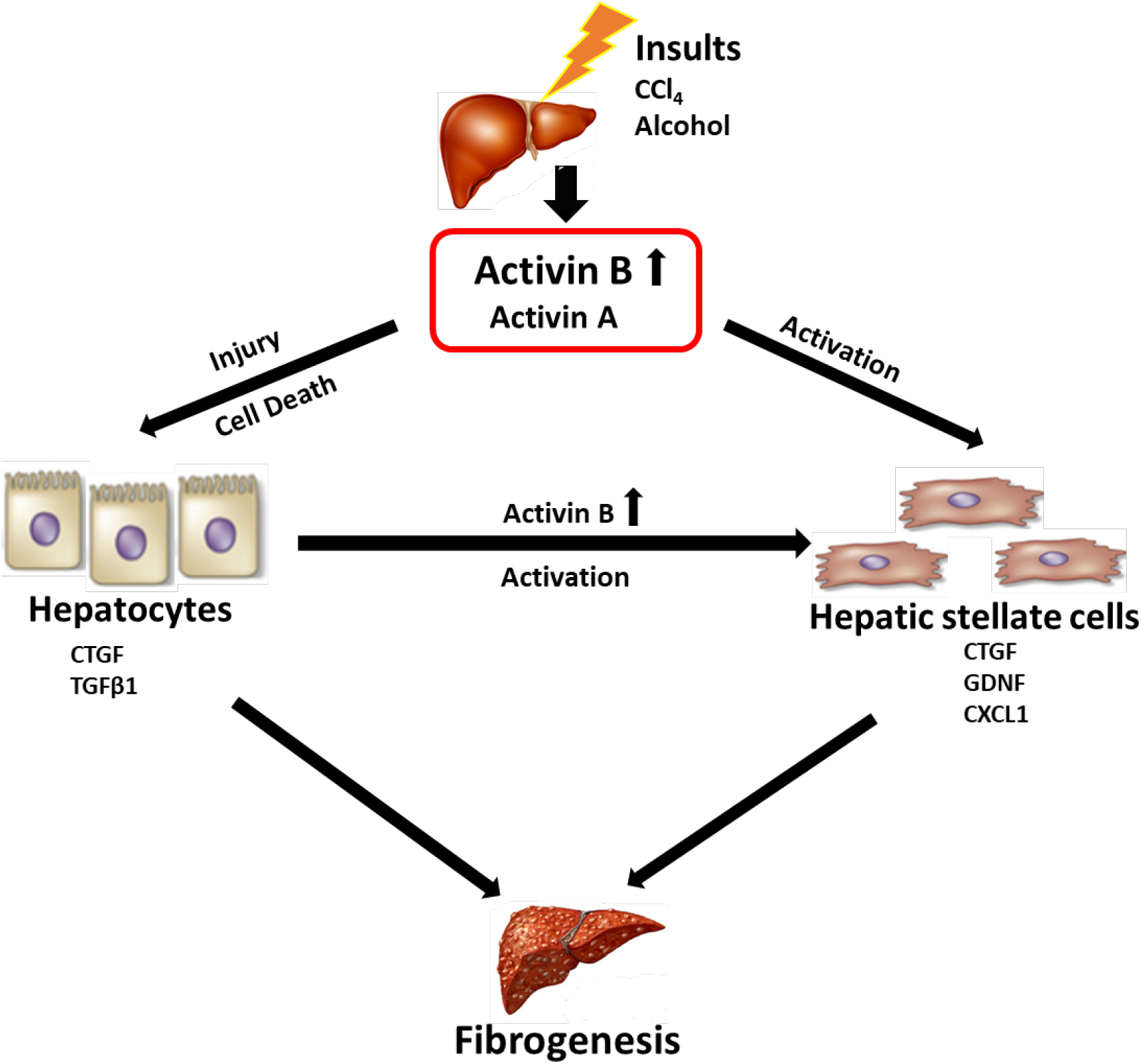
Hypothesis of the role for Activin B to modulate hepatic injury response and fibrogenesis. Hepatocytes release activin B upon injury. In an autocrine manner, activin B stimulates injured hepatocytes to produce profibrotic factors including CTGF and TGFβ1 and promotes hepatocyte death. In a paracrine manner, activin B induces hepatic stellate cells (HSCs) to produce profbrotic factors including CTGF and CXCL1 and activates HSCs to initiate liver fibrogenesis. Activin B along with other profibrotic factors collectively drive the progression of liver fibrosis. Thus, in injured liver, activin B acts as a key modulator of the initiation and progression of liver fibrosis.

## Abbreviations

TGF: transforming growth factor
CTGF: connective tissue growth factor
HSC: hepatic stellate cell
AST: aspartate aminotransferase
ALT: alanine aminotransferase
ALD: alcoholic liver disease
NASH: non-alcoholic steatohepatitis
CCl4: carbon tetrachloride
BDL: bile duct ligation
CXCL1: chemokine (C-X-C motif) ligand 1
iNOS: cytokine-inducible nitric oxide synthase

